# Integration of multiple data sources for gene network inference using genetic perturbation data

**DOI:** 10.1101/158394

**Authors:** Xiao Liang, William Chad Young, Ling-Hong Hung, Adrian E. Raftery, Ka Yee Yeung

## Abstract

**Background:** The inference of gene regulatory networks is of great interest and has various applications. The recent advances in high-throughout biological data collection have facilitated the construction and understanding of gene regulatory networks in many model organisms. However, the inference of gene networks from large-scale human genomic data can be challenging. Generally, it is difficult to identify the correct regulators for each gene in the large search space, given that the high dimensional gene expression data only provides a small number of observations for each gene.

**Results:** We present a Bayesian approach integrating external data sources with knockdown data from human cell lines to infer gene regulatory networks. In particular, we assemble multiple data sources including gene expression data, genome-wide binding data, gene ontology, known pathways and use a supervised learning framework to compute prior probabilities of regulatory relationships. We show that our integrated method improves the accuracy of inferred gene networks. We apply our method to two different human cell lines, which illustrates the general scope of our method.

**Conclusions:** We present a flexible and systematic framework for external data integration that improves the accuracy of human gene network inference while retaining efficiency. Integrating various data sources of biological information also provides a systematic way to build on knowledge from existing literature.

## Background

The inference of gene regulatory networks has attracted great interest in recent years, especially gene network inference from large-scale data. Advances in technology have led to the generation of high-throughput biological data. Gene regulatory networks play an important role in understanding the interactions between genes and have many applications. However, inferring gene networks from high-dimensional genomic data can be challenging.

We define a gene regulatory network as a directed graph that represents the regulatory relationships between genes, in which each node represents a gene and each directed edge represents the regulatory relationship between a regulator (parent node) and a target gene (child node). Furthermore, these regulatory relationships or edges from regulators to target genes can be calibrated by probabilities representing the likelihood of such edges, especially in Bayesian approaches.

There is an extensive literature on methods for the inference of human gene regulatory networks and their applications. Some authors inferred gene networks to uncover causal relationships between gene expression and disease, which could help drug discovery and development [1, 2, 3] as well as disease biomarkers [4]. Woo et *al.* [5] proposed a method to predict changes in gene expression level after drug perturbation, which offers insight into target prioritization of novel compounds. Also, gene network inference could advance the understanding of the mechanisms underlying various biological processes and identify genes that play important roles in biological activities.

Bayesian networks are one of the most commonly used modeling approaches in gene network construction. For example, Friedman et *al.* [6] built a framework on Bayesian networks to infer interactions between genes based on multiple expression measurements. A Bayesian network is a directed acyclic graph that describes the joint probabilities of the conditional independence between nodes. Bayesian network methods have been applied to yeast gene expression data and further extended using probabilistic graphical models [7]. Many other models have also been developed to infer gene regulatory networks, including ordinary differential equation methods (e.g. [8, 9]) and regression-based approaches (e.g. [10]). Ordinary differential equations have been used in both static and time-series gene expression data. These methods usually suffer from the curse of dimensionality, especially in the case of human data when the number of genes is large. Subsequently, dimension reduction techniques have been used, such as forward feature selection [11], singular value decompositions [12] and Principal Component Analysis [13].

Yeung and colleagues developed Bayesian regression-based network inference methods by integrating external data to yeast time series gene expression data [14, 15, 16, 17]. Specifically, they developed a regression framework using Bayesian Model Averaging (BMA) to select predictive variables using time series gene expression in yeast. BMA methods sample each model in the ensemble to improve the accuracy of inference. In addition, they also developed a supervised learning framework to integrate multiple data sources, including genome-wide binding data, additional gene expression data, protein-protein interaction data and prior knowledge from the literature.

Given that the high-dimensional gene expression data typically consist of limited numbers of experiments, it is difficult to identify the correct regulators for each gene in the large search space. The expected number of regulators for each target gene is relatively small compared to the whole gene set. To help the inference of gene regulatory networks, many types of external data have been used. These external data include genome-wide binding data [18], genetic interactions data [19] and protein-protein interaction data [20, 21].

## Related Work

### Time-series Gene Expression Data

Various types of data have been used to infer gene networks, including both time-series and static gene expression data [22, 23]. Compared to static data, time-series data provide much additional information from sequential time points. Using time-series gene expression data, dynamic Bayesian networks considering gene expression levels from different time points allow self-loops [24, 25, 26, 27], which are not possible in Bayesian networks due to the directed acyclic graph assumption.

Although time-series gene expression data may provide useful information from which gene regulatory relationships can be derived, it can also introduce noise and redundant information which can subsequently result in a reduction of accuracy. It is also difficult to determine the optimal number of time points profiled in the experimental design and the intervals between consecutive time points, especially after taking into account the balance of data informativeness and experiment efforts [28, 29]. Interpolation approaches using measured time points have been employed to solve the problem when the number of time points are not sufficient to infer a gene network of high accuracy. Interpolation approaches not only makes the expression levels distributed more smoothly across the time points, but also handle the estimation of time derivatives of each time point which can be utilized to build ordinary differential equations (ODE) models [30, 29]. However, interpolation cannot help to alleviate the curse of dimensionality of time-series data [31]. Due to the challenge of distinguishing signal from noise in time-series gene expression data and the increased dimensionality, gene network inference from time-series data is generally time and resource consuming.

Another limitation is that it is highly difficult to infer causality using time series gene expression data alone without additional data sources. Causality in gene regulatory networks is of great biological interest. In the context of gene networks, an inferred directed edge in the form of (*A* → *B*) means the following: gene *A* is the parent or regulator of gene *B*, and that if the expression level of gene A changes then we can expect the expression level of gene *B* will change as well. However, time-series data only provide information on statistical causality. Methods have been developed to infer directed edges by leveraging additional data sources and expert knowledge [15] or using graphical models [32]. Nevertheless, these models are limited to the inference of statistical causality.

### Perturbation Gene Expression Data

To derive directed gene regulatory networks, perturbation data such as over-expression and knockdown data has been used in many proposed methods [33, 34, 6]. As static expression data without any time points, perturbation data does not reflect any dynamic biological behavior over time but the experimental design could be used to derive a causal relationship. Specifically, after gene *A* is perturbed *(i.e.* by either knockdown or over-expression), the expression level of gene *B* is observed to change. Since the causal event (perturbation) is included in the experimental design, we can infer a directed edge (*A* → *B*). Knockdown data has been widely used in the literature in gene network inference. For example, Pinna *et al.* [35] showed the effectiveness of inferring gene networks using genetic perturbation expression data followed by graph analyses when applied to synthetic data from the DREAM4 *in silico* network challenge. Subsequently, Pinna *et al.* used non-linear ordinary differential equations (ODE) to generate additional synthetic data to optimize input parameters. Other methods applied to knockdown data include Bayesian networks [6, 36] and correlation-based Gaussian noise models [37]. In addition to accounting for causality in the experimental design, processing perturbation data is generally not too time or resource consuming.

However, the accuracy of gene network inference using knockdown data to some extent depends on the assumption that all the knocked down genes are fully suppressed in the experiments, which could potentially be difficult to accomplish in practice. The difficulties do not only come from the limitations of experimental techniques, but also from the fact that many functional genes are not able to be completely removed or knocked out, although this problem could be partially solved by partially suppressing the target gene. Perturbation experiments also require advanced lab techniques and much more resources compared to time-series and static gene expression data. Therefore, the existing data sources generally contain relatively less knockdown data. For example, in the LINCS L1000 gene expression data most of the knockdown experiments are generated using only 8 cell lines [38]. The lack of data sources could limit the usage of perturbation type data.

### Combined Data

Bonneau *et al.* proposed combining time-series and knockdown gene expression data in which a regression model coupled with bi-clustering algorithms were developed and applied to infer gene networks using data from the archaeon *Halobacterium* [39]. Other studies have combined static and knockdown gene expression data using ordinary differential equation (ODE) based methods with or without external knowledge integration [40, 41].

### Ordinary Differential Equations (ODE)

As mentioned before, many different models have been used for gene network inference including ordinary differential equations (ODE) [8, 9]. By representing the model as linear or non-linear differential equations, the interaction of expression level measured from different genes could be described by variables in the equations. Linear differential equations are generally more highly abstract in terms of describing the gene interactions and could be handled by existing linear algebra methods. Non-linear differential equations are able to describe complex behaviors with the trade-off between computational cost and strict constraints [8, 42]. Differential equations could be employed to model different types of data such as static gene expression data [43, 44] and time-series gene expression data [13].

ODEs could suffer from the curse of dimensionality given a large set of candidate genes such as human gene set. Dimension reduction techniques such as forward feature selection [11], singular value decompositions [12] and principal component analysis [13] have been used in ODE approaches to reduce dimensionality of genomic data.

### Regression-based methods

Regression models are also widely used in gene network inference. The models could be built on linear differential equations [43] or other types of functions to select the variables using a regression approach. By formulating network inference as a variable selection problem, regression-based methods could be solved using existing techniques but suffer from the curse of dimensionality. Commonly used regression-based methods include regularization methods and Bayesian Model Averaging (BMA). Regularization methods such as least absolute shrinkage and selection operator (LASSO) [45], least angle regression (LARS) [46] and elastic net [47] have been employed to model different types of data in gene network inference [48, 49, 50, 51]. Variations of BMA (Bayesian model averaging) methods have been proposed to facilitate gene network inference. Examples include iBMA [52], ScanBMA [16] and fastBMA [17] for analyzing high dimensional gene expression data.

### Bayesian Networks

Another established gene network inference approach is Bayesian networks [26, 53, 7]. A Bayesian network is a flexible framework that can be applied to both continuous and discrete types of data. It allows the incorporation of prior knowledge [54]. Although only statistical causality can be inferred by a Bayesian network, biological causality can be implied with constraints of prior knowledge or expression levels [55, 24].

However, Bayesian network inference could be time consuming given a large set of variables since it is a NP-complete problem [56, 57]. The computational complexity increases exponentially with the number of network nodes. A Bayesian network is a directed acyclic graph (DAG), hence it cannot contain any feedback loops that are ubiquitous in real-life biological systems. As mentioned before, this constraint could be removed with the application of Dynamic Bayesian networks to time-series data.

### Data Integration

Instead of using a single data source, many proposed methods have incorporated external knowledge in the construction of Bayesian networks and other models. For example, Le *et al.* [58] and Geier *et al.* [28] have applied Bayesian networks to synthetic data with prior knowledge. Imoto *et al.* [21] employed Bayesian networks on gene expression data with known regulatory interactions in yeast. Other approaches of data integration in Bayesian networks include James *et al.* [59] using gene expression data in Escherichia coli with literature knowledge, Djebbari and Quackenbush [20] integrating literature knowledge and protein-protein interaction (PPI), and Nariai *et al.* [60] integrating protein-protein interactions and pathway data. Other external knowledge integration methods include linear and non-linear differential equations [61, 62]. Data integration on human microarray data has also been conducted [61]. Multiple source knowledge integration has also been shown to lead to positive results in gene network inference [63, 64].

Integrating different sources of biological information can avoid the bias generated from a single data source. However, external knowledge can also introduce additional noise. The overall impact of data integration depends on the quality, assumptions and biases of the external knowledge.

### Our Contributions

In this paper, we present an approach integrating external data sources with knockdown data from human cell lines for predictive regulatory network inference. Our methods build on the previous work by Young and colleagues [65, 66] in which they developed a Bayesian regression framework to infer gene networks from knockdown expression data. Our key contribution is the supervised learning framework that integrates multiple data sources to derive the prior probabilities of regulatory relationships in humans. Subsequently, we incorporate these prior probabilities in the Bayesian regression-based approach to infer gene networks from knockdown data. Our results show improved accuracy of the inferred gene networks. In addition, we extend Young *et al.* [65] by applying our methods to more than one cell line (skin melanoma cell line A375 and lung cancer cell line A549).

## Methods

### LINCS L1000 Gene Expression Data

The Library of Integrated Cellular Signatures (LINCS) http://lincsproject.org [38] is a National Institutes of Health (NIH) funded program that aims to develop comprehensive signatures of cellular states and related tools. Many types of large scale data were generated to profile changes induced by genetic and drug perturbations across human cell lines. In particular, the LINCS L1000 data generated by the Broad Institute measure the gene expression levels across approximately 1,000 landmark genes. These landmark genes were chosen to capture approximately 80% of the information for 20,000 genes in the human genome. The LINCS L1000 gene expression data are publicly available from the Gene Expression Omnibus (GEO) database with accession number GSE70138 http://www.ncbi.nlm.nih.gov/geo/query/acc.cgi?acc=GSE70138.

The L1000 experiments were performed using Luminex Bead technology [67], generating high-throughput gene-expression assays using 384-well plates. To measure the expression level of specific genes, color-coded microspheres bind fluorophore and the corresponding RNA sequence. Therefore, the expression level of each gene could be represented by the intensity of the fluorescence. For a pair of genes, two types of beads sharing one bead color were designed to measure the expression of the two different genes. In each perturbation experiment, about 35,000 to 50,000 beads across 500 bead colors were added to each well to measure the expression levels of approximately 1000 landmark genes.

The beads for each pair of genes were mixed in approximately 2:1 ratio. Therefore, two peaks are expected in a histogram of fluorescence levels, and these observed peaks were deconvoluted to assign expression values to the appropriate pair of genes. To reduce the noise from experimental conditions, there are several wells used for control on each plate. In addition, technical replicate data was generated in which the same perturbations were performed in the same wells across multiple plates.

The L1000 gene expression data was generated and processed by the Broad Institute LINCS Center for Transcriptomics as part of their Connectivity Map project [68]. L1000 gene expression data is publicly available in different formats varying from levels 1 to 5 [69]. Level 1 represents the raw unprocessed data from the Luminex Bead technology. In level 2, the gene expression values of the landmark genes were deconvoluted from the observed fluorescence levels and normalized to a set of internal standards. Subsequently, quantile normalization was performed on these landmark genes, and interpolated to all 20,000 human genes in level 3. The level 4 and level 5 data consist of the gene signatures comparing the perturbed experiments to the unperturbed experiments.

Young *et al.* [66] observed that the deconvolution step introduces artifacts in the data. There are three types of artifacts found and discussed in the expression data of two paired genes on the same bead color. First, the two genes can be assigned the same expression value if their expression levels are not different enough to be distinguished. Second, together with the quantile normalization step, the deconvolution step can generate incorrect additional clusters. Finally, sometimes the paired two genes can be assigned flipped expression values, which means gene A is assigned the expression value of gene B and vice versa. A correction which can be applied to the data to eliminate some effects of these artifacts will be discussed in a later section.

As a large-scale genomics data resource, LINCS L1000 data provides a rich set of human gene expression information. Specifically, we used the knockdown experiments data in this paper. There are approximately 4500 knockdown experiments in L1000 dataset. Most of these data were generated using 8 cell lines: A375, A549, HA1E, HCC515, HEPG2, HT29, MCF7 and PC3. Data from these knockdown experiments is usually collected 96 hours after the perturbation [38].

### Method Outline

We integrate external data sources in gene network inference with human genetic perturbation data from the LINCS L1000 project. We show that the accuracy of our inferred networks is improved by external data integration and MCDC (model-based clustering with data correction). Details of the data integration and MCDC are described in the Methods section. Figure 1 shows an overview of our approach.

**Figure 1:**
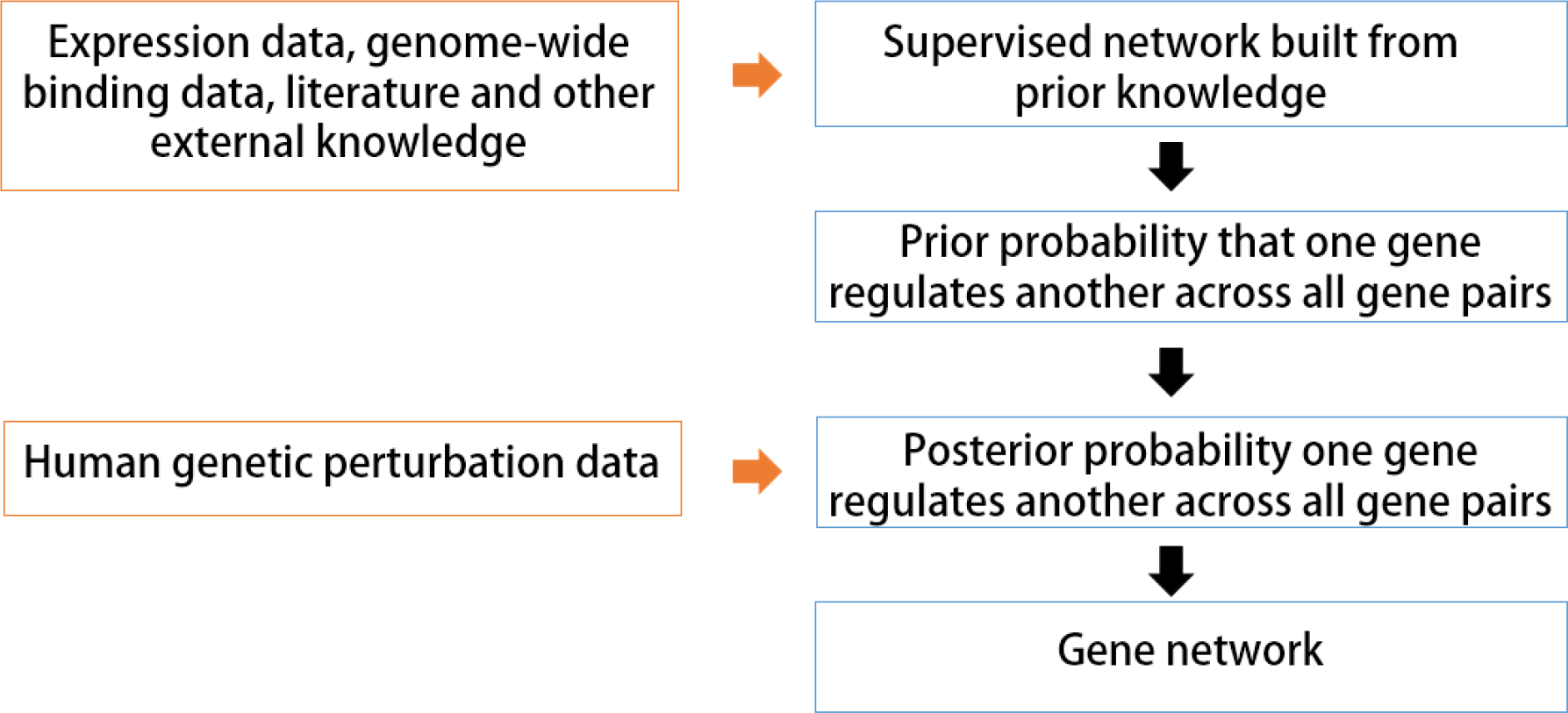
An overview of the approach. We first build a supervised framework for a selected set of target-gene regulatory pairs using external knowledge derived from the literature and existing datasets. Then, we apply machine learning methods to predict the regulatory relationships across all targetgene regulatory pairs for the landmark genes in the LINCS L1000 project. The predicted regulatory relationships are used as the prior probabilities in our Bayesian approach to predict the posterior probabilities.

Our inferred gene networks consist of directed edges in that we aim to infer causal relationship between gene pairs. This feature of our gene network does not come from the Bayesian regression-based approach but the gene knockdown experimental design. With this biological context, we assume that the observed changes in gene expression level originate from the knocked down gene.

### BayesKnockdown

We use the BayesKnockdown Bioconductor package [70] to calculate posterior probabilities of regulatory relationships. In [65], this BayesKnockdown package was applied to L1000 gene expression data from a single cell line (A375). Here we extend Young *et al.* [65] by integrating additional data sources and by applying the package to an additional cell line (A549).

To prepare the input data for the BayesKnockdown package, the LINCS L1000 knockdown experiments are first transformed by calculating z-scores to account for bias and noise among replicates:

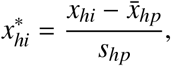

where *x*_*hi*_ represents the gene expression level of gene *h* and experiment (well) *i* on plate *p*, 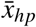 and *s*_*hp*_ are, respectively, the mean and standard deviation for gene *h* across all control experiments on plate *p*. A linear regression model is then applied to model the change in a target gene *t* as dependent on the change in the knockdown gene *h*, with 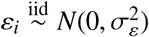 as the error term:

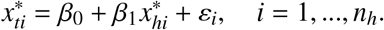

In the BayesKnockdown package [65], the linear regression model is estimated with a Bayesian approach using Zellner’s g-prior [71] for the model parameters. The parameter *g* specifies the expected size of the regression coefficient *β*_1_. The value of g can be estimated using the Expectation-Maximization algorithm [72, 16]. Then the regression model with g-prior is used to calculate the probability *Pr*(*h* → *t*|*x*) that gene *h* regulates gene *t* given the data *x*, versus the probability *Pr*_0_ that there is no regulatory relationship:

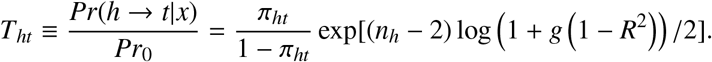

In the absence of external data sources, the prior probability of regulation *π*_ht_ is set to 0.0005 for all the gene pairs in Young *et al.* [65]. The value of 0.0005 is derived from prior knowledge for yeast data, reflecting the expected number of regulators per gene [73]. The coefficient of determination *R*^2^ for the simple linear regression is calculated from the correlation of the expression data of gene *h* and gene *t*. Then we have the posterior probability of the regulatory relationship between a given gene pair.

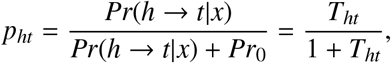

### Data integration using supervised machine learning

We download transcription factor and target gene pairs (TF-G pairs) from the PAZAR database, a public resource for transcription factor and regulatory sequence annotation [74, 75, 76]. Subsequently, we map the target genes from Ensemble IDs to Entrez IDs using Biomart [77]. After the data processing, we keep the TF-gene pairs for which both the TFs and target genes are in the L1000 landmark genes. This results in a total of 232 TF-gene pairs that we label as positive training samples (Y=1) in our supervised framework. Due to a lack of documentation on non-regulatory TF-gene pairs, we randomly generate 240 negative training samples of TF-gene pairs (Y=0) that are not documented in PAZAR.

After collecting the positive and negative training samples of TF-gene pairs, we derive the training data using external data sources to generate attributes in the supervised framework as described below. Table 1 summarizes the different types of attributes defined using external data sources in our supervised learning framework.

**Table 1:** This table summarizes the attributes in the supervised learning framework. Different versions of the same data sources with different dates (years) are considered independently. For each transcription factor gene (TF-G) pair, we have an outcome (response) value derived from the PAZAR database. For each external knowledge source, we have an attribute storing the value derived from the data source. For CCLE expression data and RNA-seq data, we calculate the correlation between the data of the paired two genes. For pathways, ChIP and Go data, we assign a binary value of 1 if the two genes appear to participate in the same biological process, otherwise a negative binary value.

| DATA SOURCE | VARIABLE | NOTES | COEFFICIENT | Pr(>\|z\|) |
| --- | --- | --- | --- | --- |
| (Intercept) | - |  | -4.072 | **0.000880** |
| CCLE | correlation | cor(TF, g) | -2.192 | **0.049747** |
| RNA-seq | correlation | cor(TF, g) | 2.043 | 0.071182 |
| BioCarta 2013 | binary | 1 if common pathway<br>0 otherwise | -0.195 | 0.927270 |
| BioCarta 2015 | binary | 1 if common pathway<br>0 otherwise | 0.705 | 0.738940 |
| KEGG 2015 | binary | 1 if common pathway<br>0 otherwise | -1.612 | 0.111851 |
| KEGG 2016 | binary | 1 if common pathway<br>0 otherwise | 0.540 | 0.342801 |
| WikiPathways 2013 | binary | 1 if common pathway<br>0 otherwise | 1.169 | 0.210120 |
| WikiPathways 2016 | binary | 1 if common pathway<br>0 otherwise | 1.506 | **0.000244** |
| Reactome 2016 | binary | 1 if common pathway<br>0 otherwise | 0.630 | **0.046281** |
| ENCODE and ChEA Consensus TFs from ChIP-X | binary | 1 if known interaction<br>0 otherwise | 1.091 | **2.02e-06** |
| ChEA 2013 | binary | 1 if known interaction<br>0 otherwise | 1.438 | **0.000690** |
| ChEA 2015 | binary | 1 if known interaction<br>0 otherwise | 1.660 | 0.192905 |
| ChIP chip | binary | 1 if known interaction<br>0 otherwise | -0.429 | 0.277970 |
| GO terms | binary | 1 if common GO term<br>0 otherwise | 0.537 | 0.141719 |

- Gene expression data across human cell lines. For each TF-gene pair, we compute the Pearson’s correlation between TF and gene across 917 human cell lines from the Cancer Cell Line Encyclopedia (CCLE) [78]. The CCLE data is publicly available from https://portals.broadinstitute.org/ccle/home and from GEO with accession number GSE36133. As another attribute (or variable) in the training data, we also compute the Pearson’s correlation between TF and gene across 675 commonly used human cell lines in the RNAseq data generated by Klijn *et al.* [79]. The Klijn *et al.* data is publicly available from ArrayExpress with accession number E-MTAB-2706 http://www.ebi.ac.uk/arrayexpress/experiments/E-MTAB-2706/.
- Gene ontology (GO). Gene Ontology (GO) defines a controlled vocabulary and descriptions of gene products across biological systems [80, 81]. Genes assigned to the same ontology terms are expected to share common functionalities. Intuitively, we expect regulatory TF-gene pairs to share common GO terms. Since GO terms are hierarchical in nature, we filter out large and hence, less informative GO terms. The upper boundary is set to 100. We define a binary attribute in our supervised framework: if a given TF-gene pair are both assigned to the same GO term, we define the binary variable to be 1, otherwise 0.
- Genome-wide binding data. We also use genome-wide binding data (ChIP-chip, ChIP-seq) from ENCODE [82]. The chromatin immunoprecipitation (ChIP) technology could be used to detect binding between proteins and DNA *in vivo.* Since transcriptional regulation is typically preceded by binding, we define a binary attribute for a (TF, gene) pair to be 1 if TF binds gene, and 0 otherwise. We derive these binary variables by parsing the processed ChIP data from the ENRICHR website [83].
- Pathways data. We hypothesize that a regulatory (TF, gene) pair is more likely to be assigned to the same biochemical pathways. Therefore, we define a binary variable for each of WikiPathways [84], KEGG [85], BioCarta [86] and Reactome [87, 88]. If TF and gene appear in the same pathway, we define the binary variable to be 1, otherwise 0. We derive these binary variables by parsing the processed library data from the ENRICHR website [83] http://amp.pharm.mssm.edu/Enrichr/.

Some attributes in our original supervised learning framework have very similar values, which decreases the informativeness of the attributes. We iteratively remove one attribute at a time and apply logistic regression to filter down to 14 attributes in our final supervised learning framework. After finalizing the training data used in the supervised learning framework, we perform 10-fold CV using different machine learning methods on our supervised framework. Table 2 summarizes the results. Logistic regression yield the highest AUROC (0.76) in our cross validation studies and is used to compute prior probabilities of regulatory relationships in next steps.

**Table 2:** This table shows the average AUROC of different machine learning models in 10 rounds of 10-fold cross validation. Models include logistic regression, SVM [89], 5-nn [90], Ada boost [91] and random forest [92]. In terms of AUROC, logistic regression and SVM are the best two models for our data.

| Model | Average AUROC | Assessment Value |
| --- | --- | --- |
| **Logistic Regression** | 0.762 | probability |
| SVM | 0.728 | probability |
| 5-nn | 0.709 | probability |
| Ada Boost | 0.669 | binary |
| Random Forest | 0.501 | binary |

### Sampling bias correction

After obtaining priors, we correct for sampling bias in the prior as previously reported in Lo *et al.* [15]. Specifically, we add an offset of *log*(π_1_/π_0_) to the log odds in our logistic regression model. Here π_1_and π_0_ are the sampling rates for positive and negative cases respectively in the training data. We use the prior knowledge from Lo *et al.* that the average number of regulators for each gene is about 2.76. Knowing there are approximately 20000 human genes, we then compute π_1_ = 232/(20000 × 2.76), π_0_ = 240/(20000 × (20000 - 2.76)) and π_1_/π_0_ = 7003.864 from our 232 positive instances and 240 negative instances in the supervised framework. Figure 2 shows the histograms before and after the correction in cell line A375. Figure 3 shows the histograms in cell line A549. Threshold probability values are set to 0.5.

**Figure 2:**
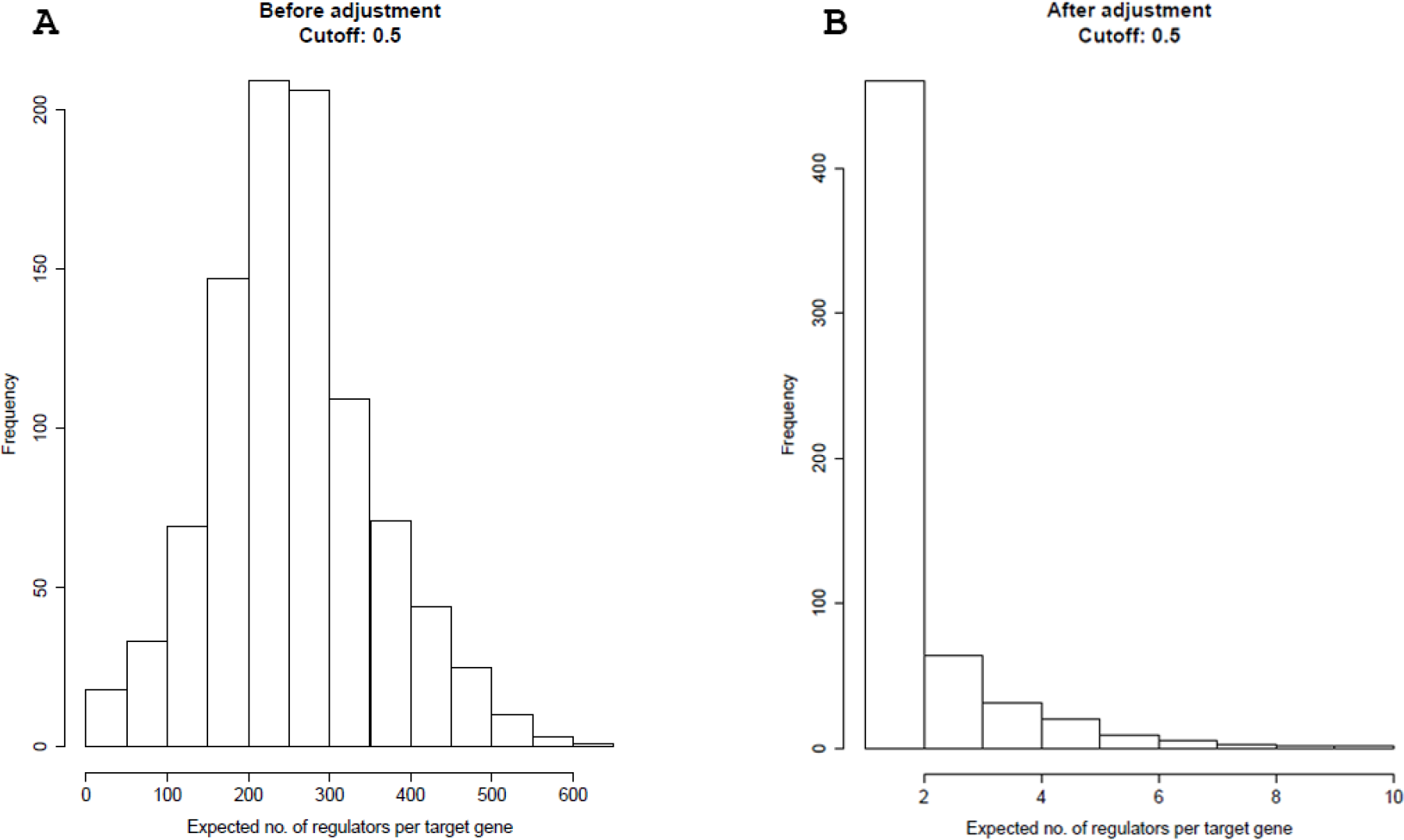
The histograms of the expected number of regulators per target gene predicted using knockdown data in cell line A375. The prior adjustment is performed only in the supervised learning step. Then the regulators are predicted using the prior probabilities combined with the L1000 knockdown data. The threshold probability value is 0.5. A) shows the histogram of the expected number of regulators per target gene without adjustment to the prior. B) shows the histogram of the expected number of regulators per target gene with adjustment to the prior.

**Figure 3:**
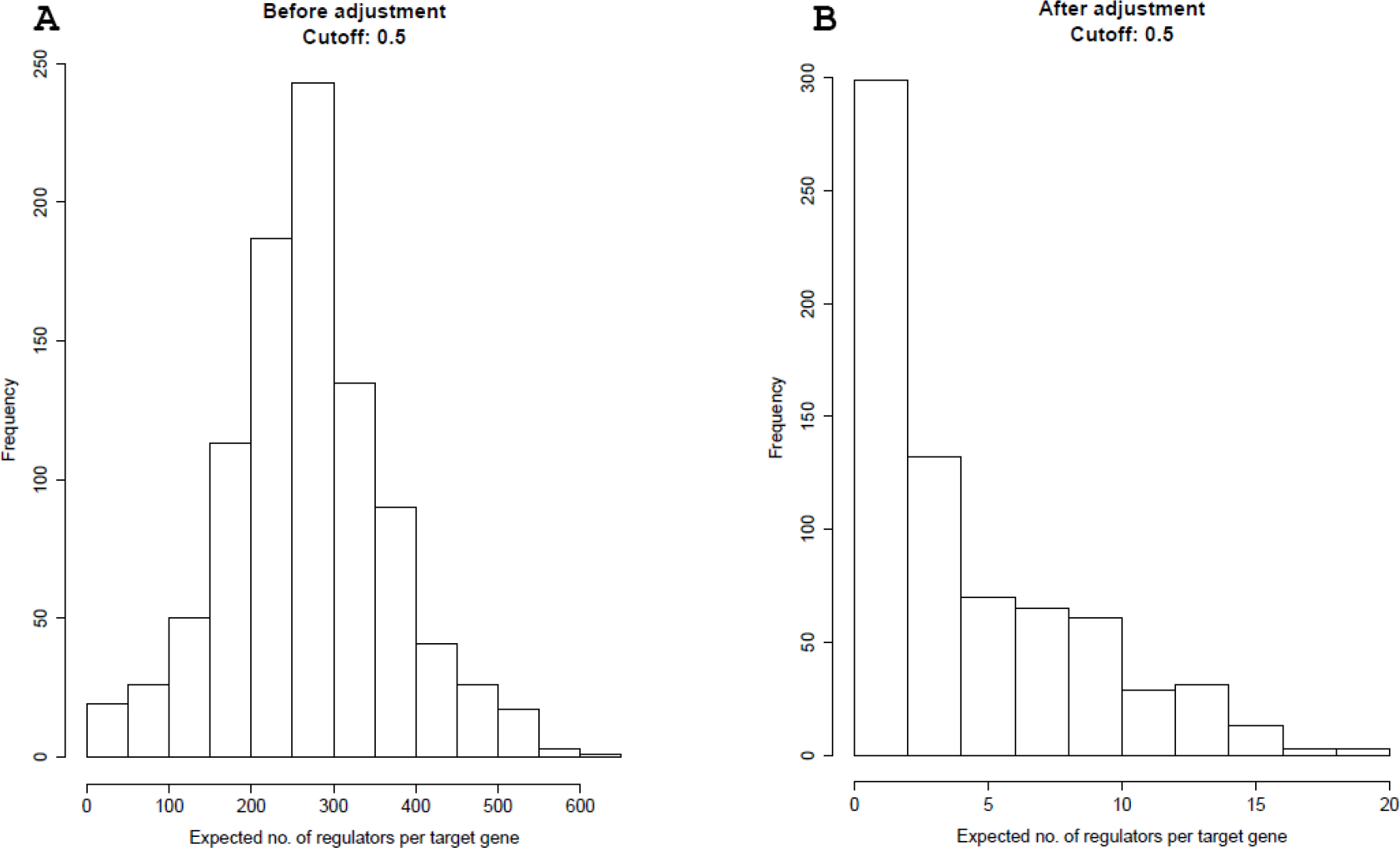
The histograms of the expected number of regulators per target gene predicted using knockdown data in cell line A549. The prior adjustment is performed only in the supervised learning step. Then the regulators are predicted using the priors combined with knockdown data. Threshold probability value is 0.5. A) shows the histogram of the expected number of regulators per target gene without adjustment to the prior. B) shows the histogram of the expected number of regulators per target gene with adjustment to the prior.

The intuition behind this correction is that our supervised training data consists of approximately the same proportion of positive and negative training samples (TF-G pairs). However, the positive cases are expected to be rare in real situations. Therefore we perform this correction to better match the biological relationships between genes in practice.

### MCDC

MCDC (Model-Based Clustering with Data Correction) is a method intended to remove artifacts in gene expression data [66]. The gene expression data was originally paired by bead color when the LINCS data was generated. In the deconvolution step during the processing, sometimes additional noisy clusters can be generated and the expression values of the paired genes can be reversed. MCDC corrects this using model-based clustering.

Model-based clustering [93, 94, 95, 96] assumes that the data come from a distribution consisting of a mixture of multiple components. Each of these components can be modeled by a Gaussian distribution with parameters which can be estimated using an EM (Expectation-Maximization) algorithm. MCDC extends model-based clustering to detect flipped points in the gene expression data.

Instead of the original data, MCDC uses a transformation matrix to determine whether each data point should be corrected. Clustering is then done with both the original and transformed data, resulting in a probability of transformation for each data point. This method thus could be used to identify flipped data points. Furthermore, to eliminate the effect of noisy clusters generated from expression level estimating process, the expression levels of the paired genes are estimated as the mean of the largest cluster after selecting the best model in MCDC.

MCDC runs with the number of clusters ranging from 1 to some maximum number. Here, the maximum number is set to 9. The best number of clusters and the best model are then selected using the BIC (Bayesian information criterion) values [96].

Our work applies MCDC to the untreated data in cell line A375 and A549. The untreated data is then used as the control to the knockdown data.

### Assessment

The high-quality transcription factor binding profile (JASPAR) database [97] provides experimentally defined transcription factor (TF) DNA-binding sites for eukaryotes. We use the TF-gene targeting relationships in this data to assess the resulting gene networks. We use Fisher’s exact test to calculate the p-values of contingency table.

A Pearson’s chi-square test is applied to a 2 × 2 contingency table to assess the consistency of our constructed network with the known regulatory relationships. Table 3 shows an example contingency table with the definitions of TP, FP, TN and FN. Precision is defined as *T P*/(*TP* + *FP*).

**Table 3:** Definition of contingency.

|  |  | Edge in T&J |  |
| --- | --- | --- | --- |
|  |  | Yes | No |
| Edge in Our Inferred network | Yes | TP | FP |
|  | No | FN | TN |

To assess our result, we use TRANSFAC and JASPAR [97] lists of edges as our reference standard. The TRANSFAC and JASPAR (T&J) edgelist contains approximately 4200 edges for 37 transcription factors that overlaps LINCS landmark genes. This is the same gold standard which was used in Young *et al.* [65, 66]. Although the T&J edgelist is limited to transcription factors that are previously well-studied, it is difficult to find a comprehensive standard for gene network assessment in mammalian systems at current stage.

## Results and Discussion

### Results:NIH LINCS Data A375

We evaluate the performance of our proposed method by comparing our inferred networks to the T&J dataset using contingency tables. Table 4 shows the assessment. We used two cutoff posterior probability values, 0.5 and 0.95, as thresholds for positive edges. The two tables in the first row are computed using knockdown data only. The two tables in the second row correspond to the network inferred using knockdown data and our external knowledge integration. The two tables in the last row show the assessment results of the network inferred from knockdown data using untreated data with MCDC correction as control, as well as external knowledge integration.

**Table 4:**
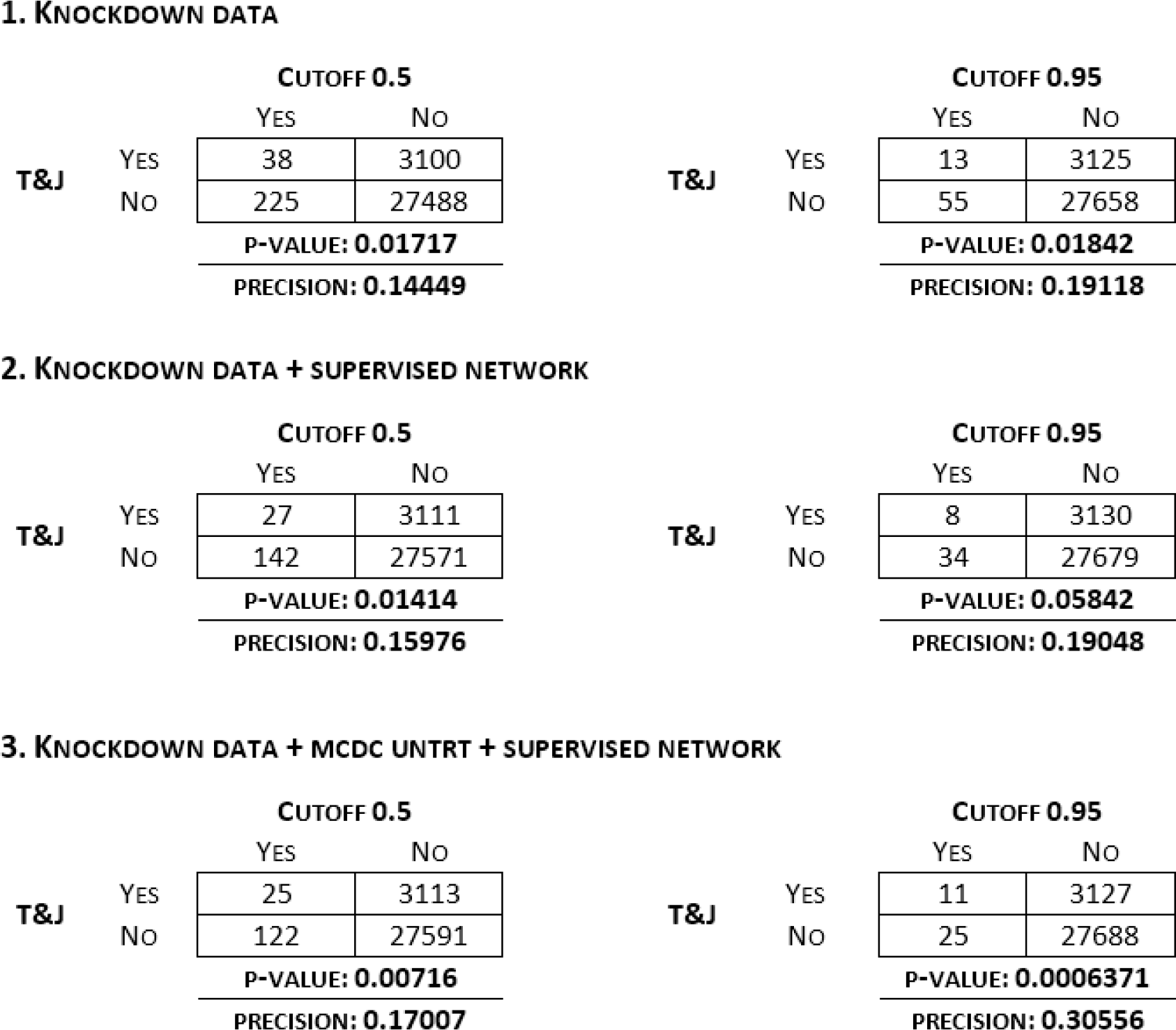
Assessment results comparing our inferred networks to TRANSFAC and JASPAR. The contingency tables display the comparison results for cell line A375 before and after external knowledge integration at 0.5 and 0.95 cutoffs. MCDC correction and external knowledge integration helps improving both the p-value and precision.

1. KNOCKDOWN DATA
|  |  | CUTOFF 0.5 |  |
| --- | --- | --- | --- |
|  |  | YES | NO |
| T&J | YES | 38 | 3100 |
|  | NO | 225 | 27488 |
|  |  | P-VALUE: **0.01717** |  |
|  |  | PRECISION: **0.14449** |  |

|  |  | CUTOFF 0.95 |  |
| --- | --- | --- | --- |
|  |  | YES | NO |
| T&J | YES | 13 | 3125 |
|  | NO | 55 | 27658 |
|  |  | P-VALUE: **0.01842** |  |
|  |  | PRECISION: **0.19118** |  |

2. KNOCKDOWN DATA + SUPERVISED NETWORK
|  |  | CUTOFF 0.5 |  |
| --- | --- | --- | --- |
|  |  | YES | NO |
| T&J | YES | 27 | 3111 |
|  | NO | 142 | 27571 |
|  |  | P-VALUE: **0.01414** |  |
|  |  | PRECISION: **0.15976** |  |

|  |  | CUTOFF 0.95 |  |
| --- | --- | --- | --- |
|  |  | YES | NO |
| T&J | YES | 8 | 3130 |
|  | NO | 34 | 27679 |
|  |  | P-VALUE: **0.05842** |  |
|  |  | PRECISION: **0.19048** |  |

3. KNOCKDOWN DATA + MCDC UNTRT + SUPERVISED NETWORK
|  |  | CUTOFF 0.5 |  |
| --- | --- | --- | --- |
|  |  | YES | NO |
| T&J | YES | 25 | 3113 |
|  | NO | 122 | 27591 |
|  |  | P-VALUE: **0.00716** |  |
|  |  | PRECISION: **0.17007** |  |

Table 4: Assessment results comparing our inferred networks to TRANSFAC and JASPAR.
|  |  | CUTOFF 0.95 |  |
| --- | --- | --- | --- |
|  |  | YES | NO |
| T&J | YES | 11 | 3127 |
|  | NO | 25 | 27688 |
|  |  | P-VALUE: **0.0006371** |  |
|  |  | PRECISION: **0.30556** |  |

We observe that both MCDC and integrating prior knowledge improve both the p-value from the Fisher’s exact test and the precision. By applying both MCDC and external knowledge integration, the p-values were improved from around 0.01 to around 0.001. Also, the precision increased from 0.14 to 0.17 at 0.5 cutoff, and 0.2 to 0.3 at 0.95 cutoff.

Next, we compare our inferred edges to the T&J dataset by ranked lists. The assessment results are shown in Table 5. We first identify all the edges in the intersection of our predicted edges and T&J edges. Then we rank these found edges by posterior probabilities from our prediction in descending order. Finally, we rank all our predicted edges by posterior probabilities in descending order. For each edge also in T&J dataset, we note down the corresponding ranking in our edgelist for the same edge. With the edge ranks we not only assess our inferred networks by the found edges, but also involved the values of posterior probabilities.

**Table 5:** Comparison of the rank of the first 25 edges found and match the TRANSFAC and JASPAR edgelist in cell line A375. Edges are ranked by posterior probability. The numbers represent the rankings of true positive edges (i.e. edges found both in our gene network and T&J edgelist) among positive edges (i.e. edges found in our network). The table shows the external knowledge integration helps improving results of middle-ranked edges, which makes the result more steady.

| FOUND EDGE | KNOCKDOWN<br>DATA | SUPERVISED<br>NETWORK | KD + SUPERVISED<br>NETWORK | KD + MCDC<br>UNTRT | KD + MCDC UNTRT +<br>SUPERVISED NETWORK |
| --- | --- | --- | --- | --- | --- |
| 1 | 6 | 2 | 7 | 2 | 2 |
| 2 | 9 | 3 | 10 | 6 | 6 |
| 3 | 11 | 7 | 11 | 8 | 8 |
| 4 | 15 | 12 | 15 | 9 | 9 |
| 5 | 22 | 26 | 33 | 12 | 13 |
| 6 | 34 | 40 | 34 | 20 | 16 |
| 7 | 39 | 54 | 37 | 22 | 23 |
| 8 | 40 | 63 | 42 | 29 | 27 |
| 9 | 41 | 64 | 43 | 42 | 28 |
| 10 | 48 | 66 | 45 | 45 | 30 |
| 11 | 56 | 88 | 65 | 49 | 35 |
| 12 | 58 | 91 | 67 | 53 | 41 |
| 13 | 65 | 106 | 73 | 56 | 56 |
| 14 | 77 | 109 | 74 | 57 | 57 |
| 15 | 78 | 110 | 80 | 66 | 58 |
| 16 | 86 | 112 | 82 | 68 | 63 |
| 17 | 98 | 114 | 83 | 71 | 80 |
| 18 | 100 | 117 | 95 | 73 | 87 |
| 19 | 102 | 118 | 100 | 83 | 90 |
| 20 | 111 | 132 | 109 | 92 | 96 |
| 21 | 119 | 136 | 113 | 93 | 97 |
| 22 | 122 | 139 | 119 | 95 | 99 |
| 23 | 130 | 151 | 122 | 102 | 102 |
| 24 | 131 | 154 | 124 | 108 | 116 |
| 25 | 135 | 156 | 127 | 112 | 123 |

As an example, for the first column “Knockdown Data”, the sixth edge in our edgelist is the first edge found in T&J in Table 5. This number means that the top 5 edges in our edgelist are not found in T&J. These ranked lists can help us to determine the differences between T&J and our own edgelist.

We can see that the external knowledge integration improves the results of middle-ranked edges. As another example, In the third column that corresponds to the network inferred using both the L1000 knockdown data and external knowledge integration, we can see that the 25th found edge is ranked 127th in our edgelist. In the first column, from knockdown data only, the 25th found edge is ranked 135th in our edgelist. The larger difference in rankings indicates a larger difference between our prediction and T&J dataset.

Figure 4 shows precision-recall curves under different combinations of data and prior. After external knowledge integration, the area under the curve has been improved. Furthermore, with the MCDC correction to the untreated data the area under the curve has been further improved.

**Figure 4:**
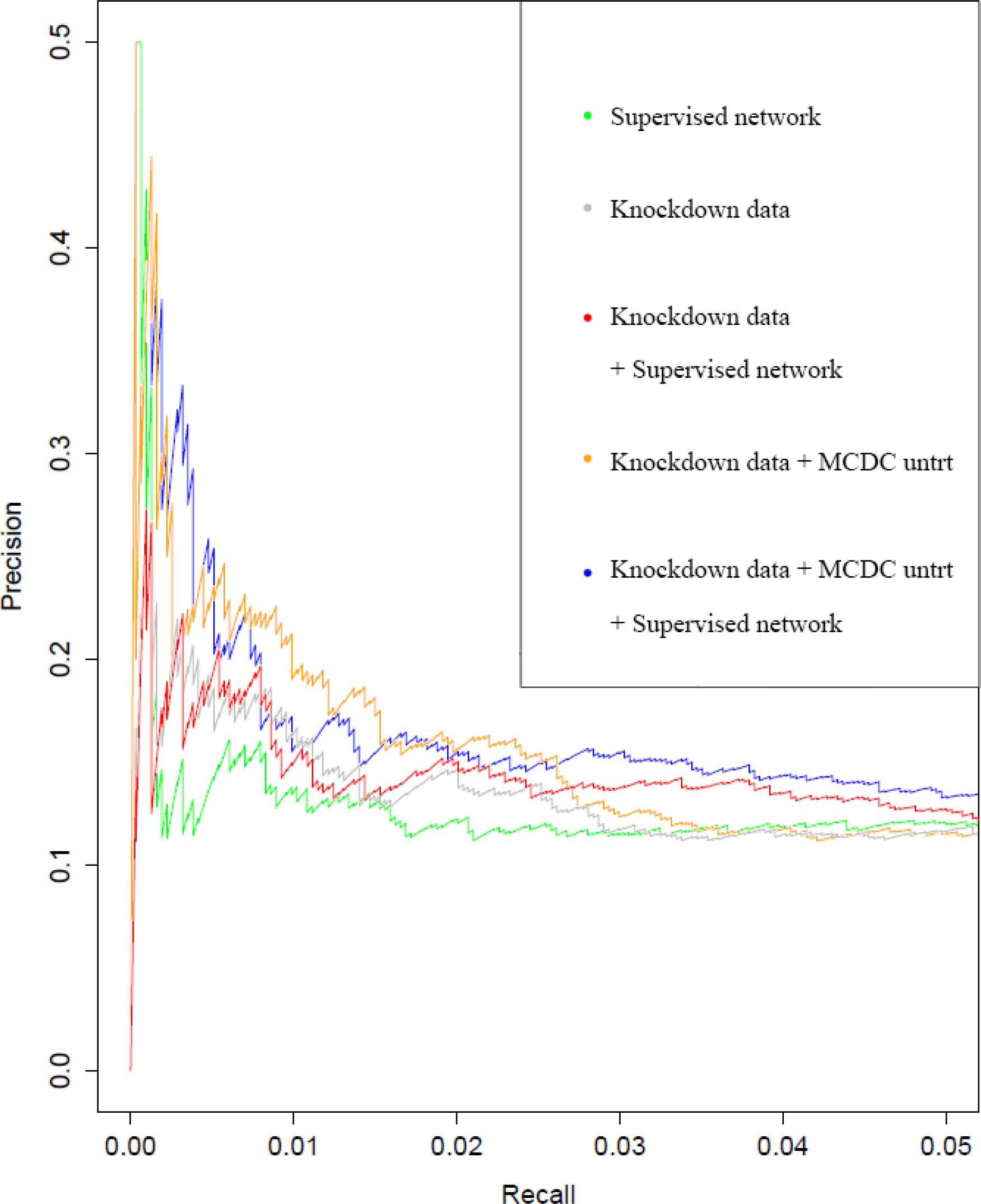
Precision-recall curves for cell line A375 using different data assessed with TRANSFAC and JASPAR. The results are improved by external knowledge integration with or without MCDC correction.

Our gene network inferred from A375 gene expression data is shown in the form of a directed graph in two figures. We used A375 knockdown data and MCDC-corrected untreated data, then integrated priors from our supervised framework to infer this gene network. Figure 5 shows all the inferred edges at a cutoff of 0.5. Figure 6 shows all the true positive edges found in TRANSFAC and JASPAR database at a cutoff of 0.5. Some of our inferred edges are also found in literature such as *CREB1 → JUN* [98, 99, 100]. Compared to another cell line we worked on, A375 cell line has less noise and therefore constructs a more reliable gene network.

**Figure 5:**
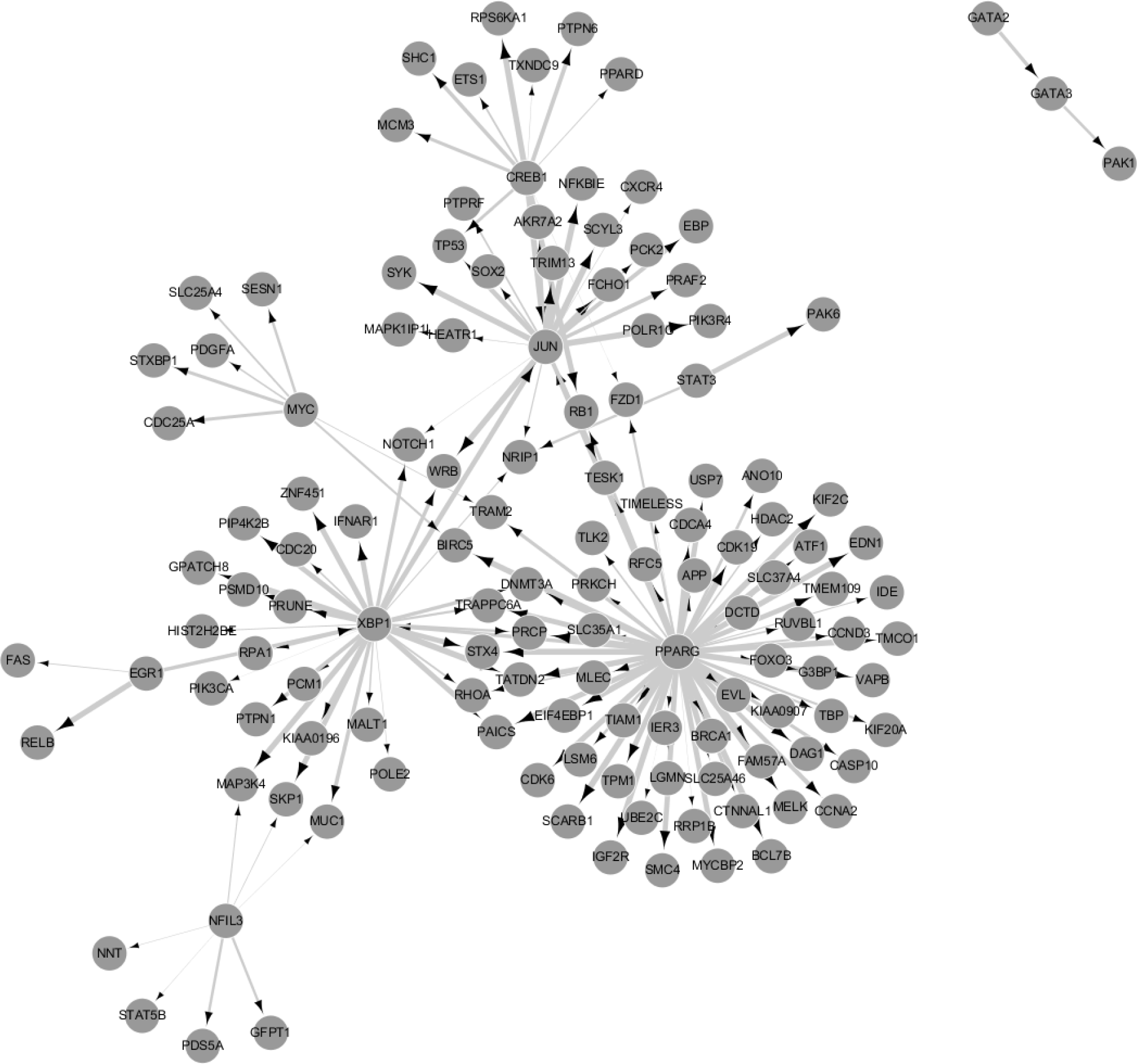
Inferred directed edges at a posterior probability cutoff of 0.5 from the gene network generated by integrating the supervised framework with the knockdown data and MCDC-corrected untreated data. Each node represents a gene and each edge represents a regulatory interaction between the two genes. The width of each edge is in proportion to the inferred posterior probability that the regulatory relationship exists for the corresponding gene pair.

**Figure 6:**
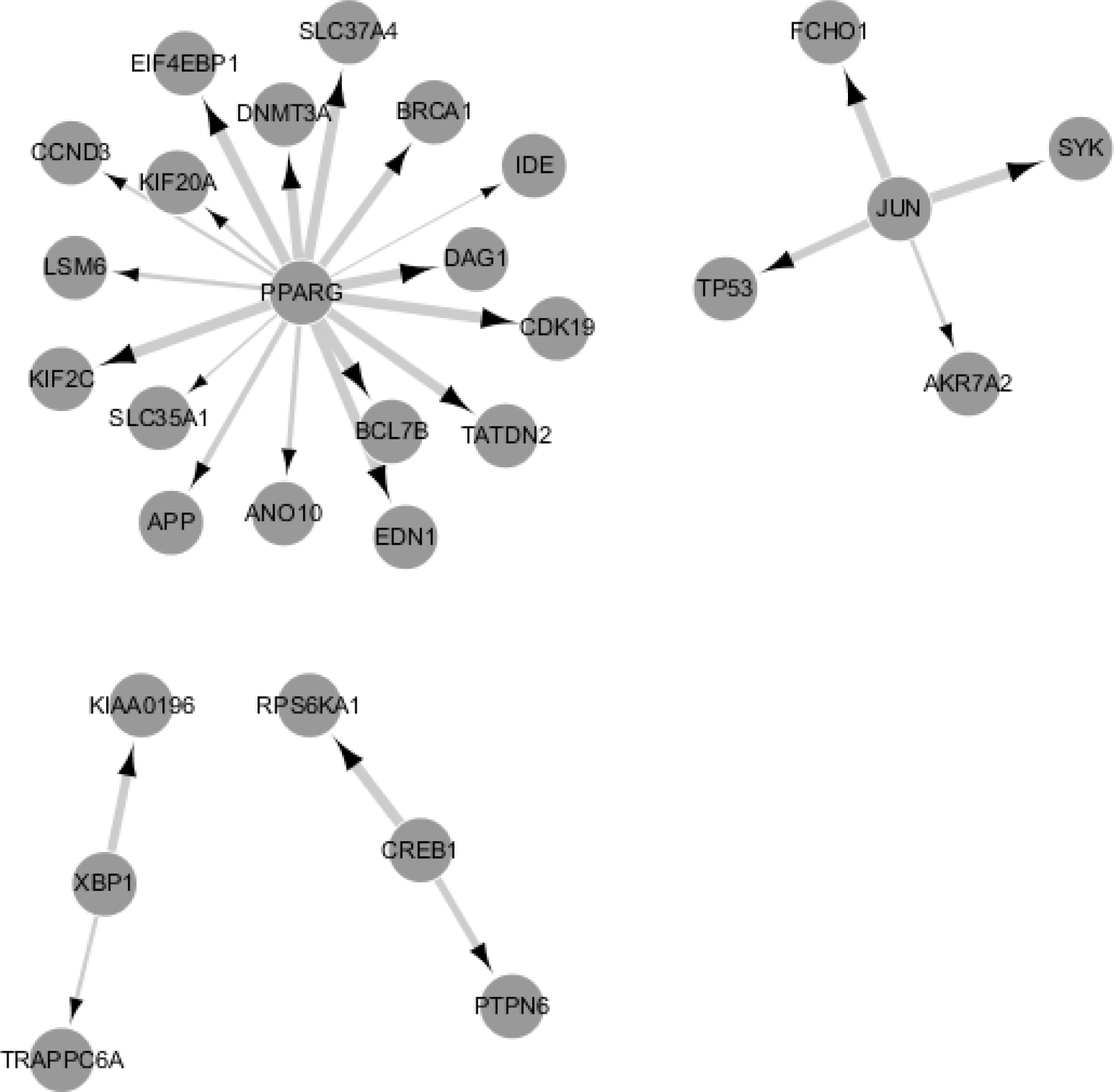
True positive edges at a posterior probability cutoff of 0.5 from the gene network generated by integrating the supervised framework with the knockdown data and MCDC-corrected untreated data. These true positive edges represent the edges from Figure 5 that are also found in our assessment criteria. Each node represents a gene and each edge represents a regulatory interaction between the two genes. The width of each edge is in proportion to the inferred posterior probability that the regulatory relationship exists for the corresponding gene pair.

### Results: Lung Cancer A549

We apply our proposed methods and assessment criteria to another cell line A549. Table 6 shows the contingency table comparing T&J and our result. The two thresholds for positive edges are again set to 0.5 and 0.95. Similar to the results on cell line A375, we observe that the MCDC correction and integrated prior knowledge also improve the p-value and precision in A549, although the effect applying MCDC is not as significant as in A375. By applying both the MCDC correction and external knowledge integration, the p-values were improved from 0.001 level to 0.0001 level. Also, the precision increased from 0.13 to 0.15 at 0.5 cutoff, and 0.14 to 0.15 at 0.95 cutoff.

**Table 6:**
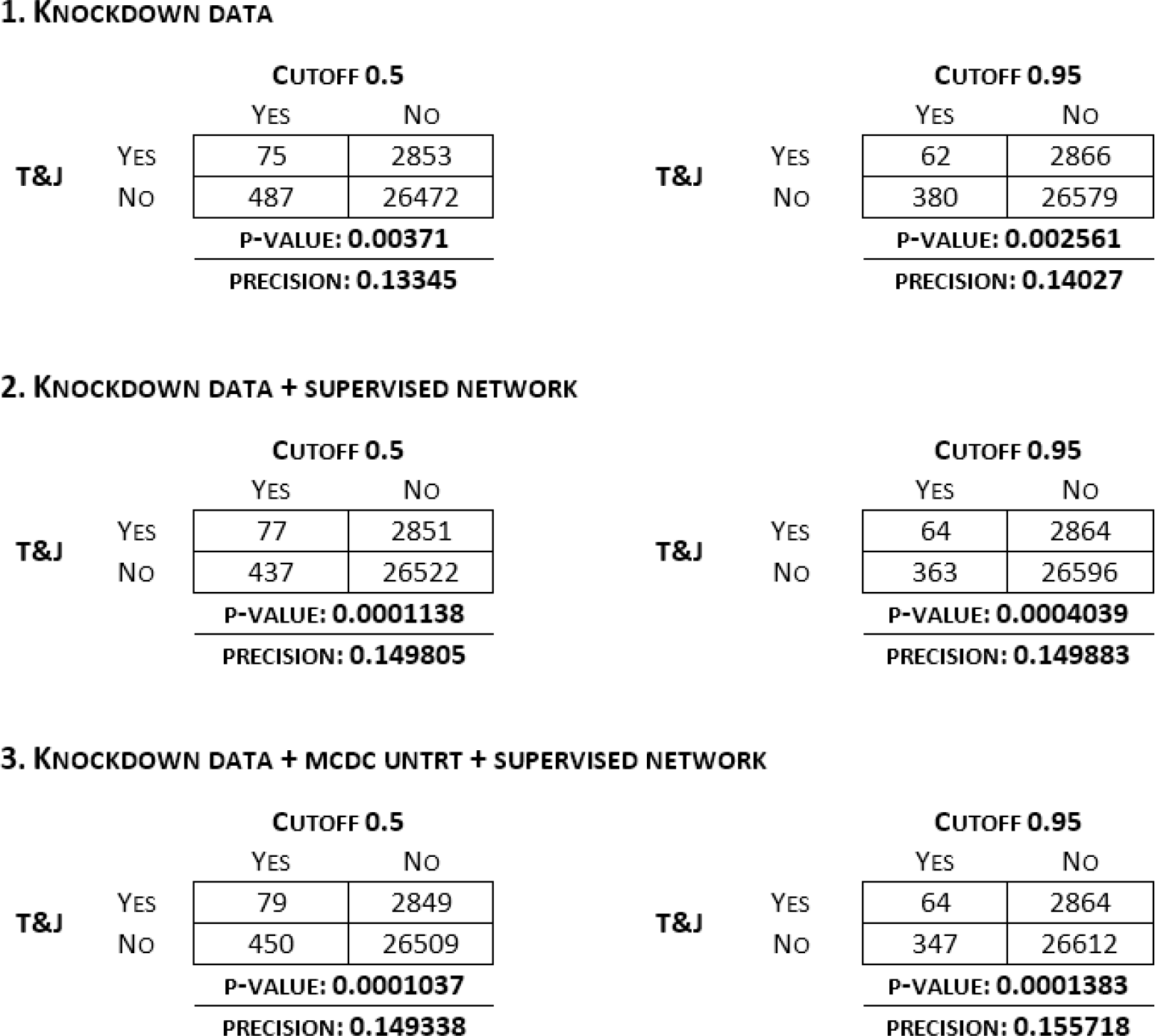
Assessment results comparing our inferred networks to TRANSFAC and JASPAR. The contingency tables display the comparison results for cell line A549 before and after external knowledge integration at 0.5 and 0.95 cutoffs. P-values have been improved by external knowledge integration at
both cutoffs.

1. KNOCKDOWN DATA
|  |  | CUTOFF 0.5 |  |
| --- | --- | --- | --- |
|  |  | YES | NO |
| T&J | YES | 75 | 2853 |
|  | NO | 487 | 26472 |
|  |  | P-VALUE: **0.00371** |  |
|  |  | PRECISION: **0.13345** |  |

|  |  | CUTOFF 0.95 |  |
| --- | --- | --- | --- |
|  |  | YES | NO |
| T&J | YES | 62 | 2866 |
|  | NO | 380 | 26579 |
|  |  | P-VALUE: **0.002561** |  |
|  |  | PRECISION: **0.14027** |  |

2. KNOCKDOWN DATA + SUPERVISED NETWORK
|  |  | CUTOFF 0.5 |  |
| --- | --- | --- | --- |
|  |  | YES | NO |
| T&J | YES | 77 | 2851 |
|  | NO | 437 | 26522 |
|  |  | P-VALUE: **0.0001138** |  |
|  |  | PRECISION: **0.149805** |  |

|  |  | CUTOFF 0.95 |  |
| --- | --- | --- | --- |
|  |  | YES | NO |
| T&J | YES | 64 | 2864 |
|  | NO | 363 | 26596 |
|  |  | P-VALUE: **0.0004039** |  |
|  |  | PRECISION: **0.149883** |  |

3. KNOCKDOWN DATA + MCDC UNTRT + SUPERVISED NETWORK
|  |  | CUTOFF 0.5 |  |
| --- | --- | --- | --- |
|  |  | YES | NO |
| T&J | YES | 79 | 2849 |
|  | NO | 450 | 26509 |
|  |  | P-VALUE: **0.0001037** |  |
|  |  | PRECISION: **0.149338** |  |

Table 6: Assessment results comparing our inferred networks to TRANSFAC and JASPAR.
|  |  | CUTOFF 0.95 |  |
| --- | --- | --- | --- |
|  |  | YES | NO |
| T&J | YES | 64 | 2864 |
|  | NO | 347 | 26612 |
|  |  | P-VALUE: **0.0001383** |  |
|  |  | PRECISION: **0.155718** |  |

The assessment results of edge ranks are shown in Table 7. As in A375, external knowledge integration and MCDC correction improve the results of middle-ranked edges. Figure 7 shows the precision-recall curves for the A549 cell line. After external knowledge integration, the area under the curve has been improved. The MCDC correction to the untreated data has not further improved the result significantly, which is different from A375.

**Table 7.** Comparison of the rank of the first 25 edges found and match the TRANSFAC and JASPAR edgelist in cell line A549. Edges are ranked by posterior probability. The numbers represent the rankings of true positive edges (i.e. edges found both in our gene network and T&J edgelist) among positive edges (i.e. edges found in our network). The table shows the external knowledge integration helps improving results of middle-ranked edges, which makes the result more steady.

| FOUND EDGE | KNOCKDOWN<br>DATA | SUPERVISED<br>NETWORK | KD + SUPERVISED<br>NETWORK | KD + MCDC<br>UNTRT | KD + MCDC UNTRT +<br>SUPERVISED NETWORK |
| --- | --- | --- | --- | --- | --- |
| 1 | 8 | 9 | 2 | 5 | 6 |
| 2 | 11 | 14 | 25 | 8 | 9 |
| 3 | 16 | 21 | 30 | 12 | 14 |
| 4 | 35 | 31 | 36 | 26 | 28 |
| 5 | 40 | 32 | 43 | 27 | 29 |
| 6 | 41 | 40 | 46 | 34 | 36 |
| 7 | 43 | 45 | 47 | 52 | 56 |
| 8 | 55 | 57 | 62 | 53 | 57 |
| 9 | 81 | 70 | 70 | 64 | 70 |
| 10 | 82 | 84 | 71 | 75 | 73 |
| 11 | 107 | 87 | 83 | 85 | 75 |
| 12 | 111 | 97 | 94 | 87 | 79 |
| 13 | 131 | 101 | 100 | 89 | 88 |
| 14 | 136 | 110 | 107 | 98 | 104 |
| 15 | 139 | 115 | 115 | 100 | 111 |
| 16 | 164 | 117 | 123 | 103 | 118 |
| 17 | 175 | 119 | 124 | 125 | 127 |
| 18 | 177 | 121 | 128 | 137 | 139 |
| 19 | 178 | 124 | 129 | 149 | 141 |
| 20 | 180 | 143 | 130 | 157 | 161 |
| 21 | 183 | 147 | 142 | 170 | 163 |
| 22 | 190 | 149 | 156 | 171 | 175 |
| 23 | 196 | 162 | 163 | 180 | 176 |
| 24 | 219 | 165 | 170 | 184 | 177 |
| 25 | 220 | 183 | 178 | 194 | 180 |

**Figure 7.**
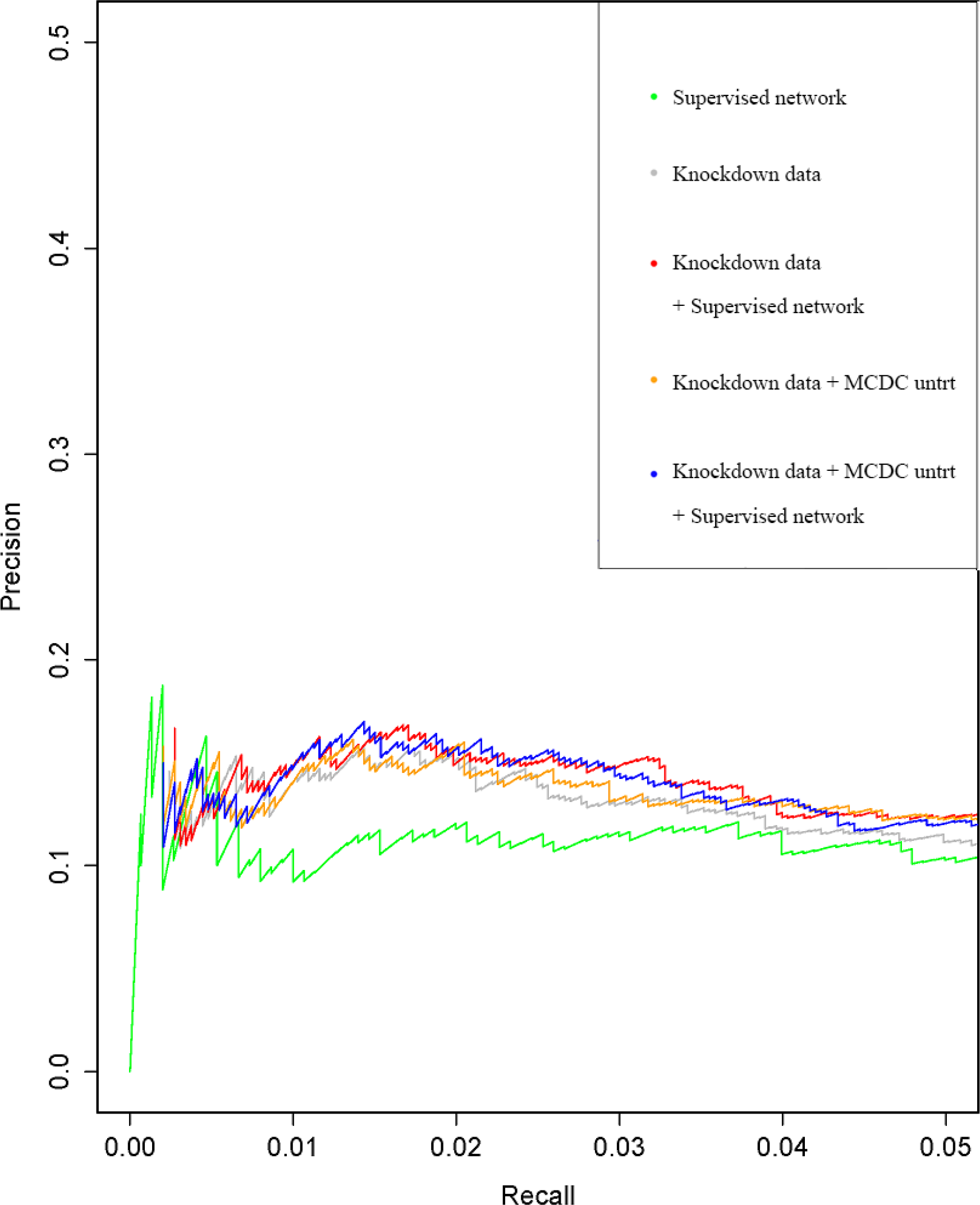
Precision-recall curves for cell line A549 using different data assessed with TRANSFAC and JASPAR. The results are improved by external knowledge integration with or without MCDC correction.

## Conclusions

In this paper, we present an approach that integrates external data sources with knockdown data from human cell lines for gene network inference. Our key contribution is to present a supervised learning framework to systematically integrate multiple data sources, and we demonstrate the flexibility of our Bayesian regression framework to integrate multiple data sources. Furthermore, the Bayesian approach we used to estimate our linear regression model is highly computationally efficient. Our ultimate aim is causal inference, and some basis for this is provided by the experimental design of the knockdown data. In addition, we extend Young *et al.* [65] by applying our methods to more than one cell line (skin melanoma cell line A375 and lung cancer cell line A549). Note that the supervised learning step is universal for all cell lines. When we apply our methods to additional cell lines, we only need to repeat the regression step.

We show that the accuracy of the inferred gene networks is improved with the help of prior knowledge integration and MCDC. One point notable from our results is that the improvement in performance varies from cell line to cell line. Studying the different response to the data integration and MCDC of different cell lines remains a meaningful future direction.

## Availability of supporting data

The scripts used in the current study are available in the GitHub repository, https://github.com/xlxlx/GeneNetworkInference

## Author’s contributions

KYY envisioned this project, identified the datasets, designed the supervised learning framework and the empirical studies. AER and WCY designed the BayesKnockDown algorithm and edited the manuscript. WCY wrote and provided the R code for BayesKnockDown and model-based clustering data correction. XL extended WCY’s R code, performed the empirical studies, compared the performance of various machine learning algorithms, and created all the figures and tables. XL and KYY wrote the manuscript. LHH participated in the design of the empirical studies, project discussions and edited the manuscript. All authors read and approved the final manuscript.

## Acknowledgements

We would like to thank Bo Ding for her contributions to the data processing and the supervised learning step. AER, LHH, KYY and WCY are supported by NIH grant U54 HL127624. AER is also supported by NIH grants R01 HD054511 and R01 HD070936.

## References

[1] Yanqing Chen et al. “Variations in DNA elucidate molecular networks that cause disease”. In: Nature 452.7186 (2008), pp. 429-435.

[2] Valur Emilsson et al. “Genetics of gene expression and its effect on disease”. In: Nature452.7186 (2008), pp. 423-428.

[3] Eric E Schadt et al. “An integrative genomics approach to infer causal associations between gene expression and disease”. In: Nature Genetics 37.7 (2005), pp. 710-717.

[4] Eric E Schadt, Alan Sachs and Stephen Friend. “Embracing complexity, inching closer to reality”. In: Sci STkE 295 (2005), p. 40.

[5] Jung Hoon Woo et al. “Elucidating compound mechanism of action by network perturbation analysis”. In: Cell 162.2 (2015), pp. 441-451.

[6] Nir Friedman et al. “Using Bayesian networks to analyze expression data”. In: Journal of Computational Biology 7. 3-4 (2000), pp. 601-620.

[7] Nir Friedman. “Inferring cellular networks using probabilistic graphical models”. In: Science303.5659 (2004), pp. 799-805.

[8] EberhardO Voit. Computational analysis of biochemical systems: a practical guide for biochemists and molecular biologists. Cambridge University Press, 2000.

[9] Mads Kffirn et al. “Stochasticity in gene expression: from theories to phenotypes”. In: Nature Reviews Genetics 6.6 (2005), pp. 451-464.

[10] Eran Segal et al. “Module networks: identifying regulatory modules and their condition-specific regulators from gene expression data”. In: Nature Genetics 34.2 (2003), pp. 166-176.

[11] Tao Huang et al. “Using GeneReg to construct time delay gene regulatory networks”. In: BMC Research Notes 3.1 (2010), p. 142.

[12] Shu-Qin Zhang et al. “A new multiple regression approach for the construction of genetic regulatory networks”. In: Artificial Intelligence in Medicine 48.2 (2010), pp. 153-160.

[13] Mukesh Bansal, Giusy Della Gatta and Diego Di Bernardo. “Inference of gene regulatory networks and compound mode of action from time course gene expression profiles”. In: Bioinformatics 22.7 (2006), pp. 815-822.

[14] Yeung Ka Yee, et al. “Construction of regulatory networks using expression time-series data of a genotyped population”. In: Proceedings of the National Academy of Sciences 108.48 (2011), pp. 19436-19441.

[15] Kenneth Lo et al. “Integrating external biological knowledge in the construction of regulatory networks from time-series expression data”. In: BMC Systems Biology 6.1 (2012), p. 1.

[16] William Chad Young, Adrian E Raftery and Ka Yee Yeung. “Fast Bayesian inference for gene regulatory networks using ScanBMA”. In: BMC Systems Biology 8.1 (2014), p. 47.

[17] Ling-Hong Hung, Kaiyuan Shi, Migao Wu, William Chad Young, Adrian E. Raftery and Ka Yee Yeung. “fastBMA: Scalable Network Inference and Transitive Reduction”. In: BioRxiv 099036 (2017).

[18] David S Johnson et al. “Genome-wide mapping of in vivo protein-DNA interactions”. In: Science 316.5830 (2007), pp. 1497-1502.

[19] Michael Costanzo et al. “The genetic landscape of a cell”. In: Science 327.5964 (2010), pp. 425-431.

[20] Amira Djebbari and John Quackenbush. “Seeded Bayesian Networks: constructing genetic networks from microarray data”. In: BMC Systems Biology 2.1 (2008), p. 1.

[21] Seiya Imoto et al. “Bayesian network and nonparametric heteroscedastic regression for nonlinear modeling of genetic network”. In: Journal of Bioinformatics and Computational Biology1.02 (2003), pp. 231-252.

[22] Patrik D’haeseleer et al. “Linear modeling of mRNA expression levels during CNS development and injury.” In: Pacific Symposium on Biocomputing. Vol. 4. 1. Citeseer. 1999, pp. 4152.

[23] Reinhard Guthke et al. “Dynamic network reconstruction from gene expression data applied to immune response during bacterial infection”. In: Bioinformatics 21.8 (2005), pp. 1626-1634.

[24] Jun Zhu et al. “Characterizing dynamic changes in the human blood transcriptional network”. In: PLoS Comput Biol 6.2 (2010), e1000671.

[25] Sun Yong Kim, Seiya Imoto and Satoru Miyano. “Inferring gene networks from time series microarray data using dynamic Bayesian networks”. In: Briefings in Bioinformatics 4.3 (2003), pp. 228-235.

[26] Kevin Murphy, Saira Mian, et al. Modelling gene expression data using dynamic Bayesian networks. Tech. rep. Technical report, Computer Science Division, University of California, Berkeley, CA, 1999.

[27] Jing Yu et al. “Advances to Bayesian network inference for generating causal networks from observational biological data”. In: Bioinformatics 20.18 (2004), pp. 3594-3603.

[28] Florian Geier, Jens Timmer and Christian Fleck. “Reconstructing gene-regulatory networks from time series, knock-out data, and prior knowledge”. In: BMC Systems Biology 1.1 (2007), p. 11.

[29] MK Stephen Yeung, Jesper Tegner, James J Collins. “Reverse engineering gene networks using singular value decomposition and robust regression”. In: Proceedings of the National Academy of Sciences 99.9 (2002), pp. 6163-6168.

[30] Ting Chen, Hongyu L He, George M Church, et al. “Modeling gene expression with differential equations.” In: Pacific symposium on biocomputing. Vol. 4. 29. 1999, p. 40.

[31] Lodewyk FA Wessels, Eugene P van Someren, Marcel JT Reinders, et al. “A comparison of genetic network models.” In: Pacific Symposium on Biocomputing. Vol. 6. 4. 2001, pp. 508-519.

[32] Judea Pearl. Probabilistic reasoning in intelligent systems: networks of plausible inference. Morgan Kaufmann, 2014.

[33] Andrew Fire et al. “Potent and specific genetic interference by double-stranded RNA in Caenorhabditis elegans”. In: Nature 391.6669 (1998), pp. 806-811.

[34] Craig C Mello and Darryl Conte. “Revealing the world of RNA interference”. In: Nature431.7006 (2004), pp. 338-342.

[35] Andrea Pinna, Nicola Soranzo and Alberto De La Fuente. “From knockouts to networks: establishing direct cause-effect relationships through graph analysis”. In: PloS One 5.10 (2010), e12912.

[36] Simon Rogers and Mark Girolami. “A Bayesian regression approach to the inference of regulatory networks from gene expression data”. In: Bioinformatics 21.14 (2005), pp. 3131-3137.

[37] Faridah Hani Mohamed Salleh et al. “Reconstructing gene regulatory networks from knockout data using Gaussian Noise Model and Pearson Correlation Coefficient”. In: Computational Biology and Chemistry 59 (2015), pp. 3-14.

[38] Qiaonan Duan et al. “LINCS Canvas Browser: interactive web app to query, browse and interrogate LINCS L1000 gene expression signatures”. In: Nucleic Acids Research (2014), gku476.

[39] Richard Bonneau et al. “The Inferelator: an algorithm for learning parsimonious regulatory networks from systems-biology data sets de novo”. In: Genome Biology 7.5 (2006), R36.

[40] Ali Shojaie et al. “Inferring regulatory networks by combining perturbation screens and steady state gene expression profiles”. In: PloS One 9.2 (2014), e82393.

[41] Scott Christley, Qing Nie and Xiaohui Xie. “Incorporating existing network information into gene network inference”. In: PloS One 4.8 (2009), e6799.

[42] Hidde De Jong. “Modeling and simulation of genetic regulatory systems: a literature review”. In: Journal of Computational Biology 9.1 (2002), pp. 67-103.

[43] Timothy S Gardner et al. “Inferring genetic networks and identifying compound mode of action via expression profiling”. In: Science 301.5629 (2003), pp. 102-105.

[44] Diego di Bernardo et al. “Chemogenomic profiling on a genome-wide scale using reverse-engineered gene networks”. In: Nature Biotechnology 23.3 (2005), pp. 377-383.

[45] Robert Tibshirani. “Regression shrinkage and selection via the lasso”. In: Journal of the Royal Statistical Society. Series B (Methodological) (1996), pp. 267-288.

[46] Bradley Efron et al. “Least angle regression”. In: The Annals of Statistics 32.2 (2004), pp. 407499.

[47] Hui Zou and Trevor Hastie. “Regularization and variable selection via the elastic net”. In: Journal of the Royal Statistical Society: Series B (Statistical Methodology) 67.2 (2005), pp. 301-320.

[48] Camille Charbonnier, Julien Chiquet and Christophe Ambroise. “Weighted-LASSO for structured network inference from time course data”. In: Statistical Applications in Genetics and Molecular Biology 9.1 (2010), p. 15.

[49] Eugene P van Someren et al. “Least absolute regression network analysis of the murine osteoblast differentiation network”. In: Bioinformatics 22.4 (2006), pp. 477-484.

[50] Mika Gustafsson and Michael Hörnquist. “Gene expression prediction by soft integration and the Elastic Net—Best performance of the DREAM3 gene expression challenge”. In: PLoS One 5.2 (2010), e9134.

[51] Jie Peng et al. “Regularized multivariate regression for identifying master predictors with application to integrative genomics study of breast cancer”. In: The Annals of Applied Statistics4.1 (2010), p. 53.

[52] Ka Yee Yeung, Roger E Bumgarner and Adrian E Raftery. “Bayesian model averaging: development of an improved multi-class, gene selection and classification tool for microarray data”. In: Bioinformatics 21.10 (2005), pp. 2394-2402.

[53] Thomas Dyhre Nielsen and Finn Verner Jensen. Bayesian networks and decision graphs. Springer Science & Business Media, 2009.

[54] Adriano V Werhli, Dirk Husmeier, et al. “Reconstructing gene regulatory networks with Bayesian networks by combining expression data with multiple sources of prior knowledge”. In: Stat Appl Genet Mol Biol 6.1 (2007), p. 15.

[55] Eric E Schadt. “Molecular networks as sensors and drivers of common human diseases”. In: Nature 461.7261 (2009), pp. 218-223.

[56] David Maxwell Chickering. “Learning Bayesian networks is NP-complete”. In: Learning From Data. Springer, 1996, pp. 121-130.

[57] David Maxwell Chickering, David Heckerman and Christopher Meek. “Large-sample learning of Bayesian networks is NP-hard”. In: Journal of Machine Learning Research 5.Oct (2004), pp. 1287-1330.

[58] Phillip P Le, Amit Bahl and Lyle H Ungar. “Using prior knowledge to improve genetic network reconstruction from microarray data”. In: In Silico Biology 4.3 (2004), pp. 335-353.

[59] Gareth M James et al. “Sparse regulatory networks”. In: The Annals of Applied Statistics 4.2 (2010), p. 663.

[60] N Nariai et al. “Using protein-protein interactions for refining gene networks estimated from microarray data by Bayesian networks”. In: Pacific Symposium on Biocomputing (PSB03). 2003, pp. 336-347.

[61] Dirk Koczan et al. “Molecular discrimination of responders and nonresponders to anti-TNFalpha therapy in rheumatoid arthritis by etanercept”. In: Arthritis Research & Therapy 10.3 (2008), R50.

[62] Christian Spieth et al. “Inferring Regulatory Systems with Noisy Pathway Information.” In: German Conference on Bioinformatics. Citeseer. 2005, pp. 193-203.

[63] Jun Zhu et al. “Integrating large-scale functional genomic data to dissect the complexity of yeast regulatory networks”. In: Nature Genetics 40.7 (2008), pp. 854-861.

[64] Kevin Y Yip et al. “Improved reconstruction of in silico gene regulatory networks by integrating knockout and perturbation data”. In: PloS One 5.1 (2010), e8121.

[65] William Chad Young, Adrian E. Raftery and Ka Yee Yeung. “A Posterior Probability Approach for Gene Regulatory Network Inference in Genetic Perturbation Data”. In: Mathematical Biosciences and Engineering 13.6 (2016), pp. 1241-1251.

[66] William Chad Young, Ka Yee Yeung and Adrian E Raftery. “Model-based clustering with data correction for removing artifacts in gene expression data”. In: arXiv preprint arXiv:1602.06316(2016).

[67] Sherry A Dunbar. “Applications of Luminex0 xMAPTM technology for rapid, high-throughput multiplexed nucleic acid detection”. In: Clinica Chimica Acta 363.1 (2006), pp. 71-82.

[68] Justin Lamb et al. “The Connectivity Map: using gene-expression signatures to connect small molecules, genes, and disease”. In: Science 313.5795 (2006), pp. 1929-1935.

[69] LINCS Workflow: L1000 data processing. http://lincsproject.org/LINCS/tools/workflows/find-the-best-place-to-obtain-the-lines-11000-data. Last accessed April, 2017.

[70] BayesKnockdown package. https://bioconductor.org/packages/release/bioc/html/BayesKnockdown.html. Last accessed February, 2017.

[71] Arnold Zellner. “On assessing prior distributions and Bayesian regression analysis with g-prior distributions”. In: Bayesian Inference and Decision Techniques: Essays in Honor of Bruno De Finetti 6 (1986), pp. 233-243.

[72] Arthur P Dempster, Nan M Laird and Donald B Rubin. “Maximum likelihood from incomplete data via the EM algorithm”. In: Journal of the Royal Statistical Society. Series B (methodological) (1977), pp. 1-38.

[73] Nabil Guelzim et al. “Topological and causal structure of the yeast transcriptional regulatory network”. In: Nature Genetics 31.1 (2002), pp. 60-63.

[74] Elodie Portales-Casamar et al. “PAZAR: a framework for collection and dissemination of cis-regulatory sequence annotation”. In: Genome Biology 8.10 (2007), R207.

[75] Elodie Portales-Casamar et al. “The PAZAR database of gene regulatory information coupled to the ORCA toolkit for the study of regulatory sequences”. In: Nucleic Acids Research37.suppl 1 (2009), pp. D54-D60.

[76] PAZAR, public database of transcription factors and regulatory sequence annotation. http://www.pazar.info/. Last accessed February, 2017.

[77] BioMart. http://www.biomart.org/. Last accessedFebruary, 2017.

[78] Jordi Barretina et al. “The Cancer Cell Line Encyclopedia enables predictive modelling of anticancer drug sensitivity”. In: Nature 483.7391 (2012), pp. 603-607.

[79] Christiaan Klijn et al. “A comprehensive transcriptional portrait of human cancer cell lines”. In: Nature Biotechnology 33.3 (2015), pp. 306-312.

[80] Michael Ashburner et al. “Gene Ontology: tool for the unification of biology”. In: Nature Genetics 25.1 (2000), pp. 25-29.

[81] Gene Ontology Consortium et al. “Gene ontology consortium: going forward”. In: Nucleic Acids Research 43.D1 (2015), pp. D1049-D1056.

[82] ENCODE Project Consortium et al. “An integrated encyclopedia of DNA elements in the human genome”. In: Nature 489.7414 (2012), pp. 57-74.

[83] Edward Y Chen et al. “Enrichr: interactive and collaborative HTML5 gene list enrichment analysis tool”. In: BMC Bioinformatics 14.1 (2013), p. 128.

[84] Thomas Kelder et al. “WikiPathways: building research communities on biological pathways”. In: Nucleic Acids Research 40.D1 (2012), pp. D1301-D1307.

[85] Minoru Kanehisa and Susumu Goto. “KEGG: kyoto encyclopedia of genes and genomes”. In: Nucleic Acids Research 28.1 (2000), pp. 27-30.

[86] Darryl Nishimura. “BioCarta”. In: Biotech Software & Internet Report: The Computer Software Journal for Scient 2.3 (2001), pp. 117-120.

[87] David Croft et al. “The Reactome pathway knowledgebase”. In: Nucleic Acids Research 42.D1 (2014), pp. D472-D477.

[88] Antonio Fabregat et al. “The reactome pathway knowledgebase”. In: Nucleic Acids Research 44.D1 (2016), pp. D481-D487.

[89] e1071 package. https://cran.r-project.org/package=e1071. Last accessedFebruary, 2017.

[90] class package. https://cran.r-project.org/package=class. Last accessedFebruary, 2017.

[91] ada package. https://cran.r-project.org/package=ada. Last accessedFebruary, 2017.

[92] randomForestpackage. https://cran.r-project.org/package=randomForest. Last accessedFebruary, 2017.

[93] John H Wolfe. “Pattern clustering by multivariate mixture analysis”. In: Multivariate Behavioral Research 5.3 (1970), pp. 329-350.

[94] Jeffrey D Banfield Adrian E Raftery. “Model-based Gaussian and non-Gaussian clustering”. In: Biometrics (1993), pp. 803-821.

[95] Geoffrey McLachlan and David Peel. Finite mixture models. John Wiley & Sons, 2004.

[96] Chris Fraley and Adrian E SRafteryachs. “Model-based clustering, discriminant analysis, and density estimation”. In: Journal of the American Statistical Association 97.458 (2002), pp. 611-631.

[97] Anthony Mathelier et al. “JASPAR 2014: an extensively expanded and updated open-access database of transcription factor binding profiles”. In: Nucleic Acids Research 42.D1 (2013), pp. D142-D147.

[98] Doris M Benbrook and Nicholas C Jones. “Heterodimer formation between CREB and JUN proteins”. In: Oncogene 5.3 (1990), pp. 295-302.

[99] Denise J Spring and Edwin G Krebs. “Deletion of 11 Amino Acids in p90 rsk-mo-1Abolishes Kinase Activity”. In: Molecular and Cellular Biology 19.1 (1999), pp. 317-320.

[100] Nian Liu et al. “Unique regulation of immediate early gene and tyrosine hydroxylase expression in the odor-deprived mouse olfactory bulb”. In: Journal of Biological Chemistry 274.5 (1999), pp. 3042-3047.

